# Wnt Signalling Controls the Response to Mechanical Loading during Zebrafish Joint Development

**DOI:** 10.1101/115105

**Authors:** L H Brunt, K Begg, E Kague, S Cross, C L Hammond

## Abstract

Joint morphogenesis requires mechanical activity during development. Loss of mechanical strain causes abnormal joint development, which can impact long term joint health. While cell orientation and proliferation are known to shape the joint, dynamic imaging of developing joints *in vivo* have not been possible in other species. Using genetic labelling techniques in zebrafish we were able, for the first time, to dynamically track cell behaviours in intact moving joints. We identify that proliferation and migration, which contribute to joint morphogenesis, are mechanically controlled and are significantly reduced in immobilised larvae. By comparison to strain maps of the developing skeleton we identify canonical Wnt signalling as a candidate to transduce mechanical forces into joint cell behaviours. We show that in the jaw Wnt signalling is reduced specifically in regions of high strain in response to loss of muscle activity. By pharmacological manipulation of canonical Wnt signalling we demonstrate that Wnt acts downstream of mechanical activity and is required for joint patterning and chondrocyte maturation. Wnt16, independent of muscle activity, controls proliferation and migration, but plays no role in chondrocyte intercalation.

## Introduction

The developing skeleton is subject to many biomechanical forces, including those from fetal/early postnatal muscle activity. It has become clear from studies on animal models that mechanical stimuli are required for accurate functional joint formation (Nowlan et al., 2010b; Rolfe et al., 2013). For example, *Pax3^sp/sp^*(*splotch*) mutant mice, that lack limb muscle and *Myf5^-/-^ MyoD^-/-^* double mutants that lack all muscle, show altered morphology in many joints including elbows and shoulders (Gomez et al., 2007; Kahn et al., 2009; Nowlan et al., 2010a; Rot-Nikcevic et al., 2007; Rot-Nikcevic et al., 2006). In chick, paralysis or removal of muscle through grafts results in a knee joint that lacks refinement (Murray and Selby, 1930; Roddy et al., 2009). Zebrafish mutants that lack neuromuscular nicotinic receptors (*nic* b107) and are therefore immobile, display jaw morphology abnormalities, such as smaller and wider elements (Shwartz et al., 2012). Zebrafish jaw joint morphology is also affected by paralysis, particularly in regions associated with high compressive strain (Brunt et al., 2015; Brunt et al., 2016b). In humans, a biomechanical stimulus in utero and in newborns has a longterm impact on skeletal health (Reviewed in Shea et al., 2015). For example, Foetal Akinesia Deformation Sequence, (FADS), can cause arthrogryposis due to reduced foetal movement (Nayak et al., 2014). Risk factors such as breech birth (Luterkort et al., 1986) and swaddling that restrict hip joint movement (Clarke, 2014) can cause Developmental Dysplasia of the hip (DDH), (Sugano et al., 1998). If left untreated, the abnormal joint shape in DDH can lead to early onset osteoarthritis (OA) (Mavcic et al., 2008). Thus, diverse vertebrate species ranging from fish to humans rely on muscle activity to provide mechanical stimuli to activate the cellular processes required to shape joints during development. This process also has an impact on joint function and health later in life.

Mechanical stimulus can activate genes important for skeletogenesis. *In vitro* experiments have shown that application of force to chondrocytes can lead to activation of genes, including those encoding cartilage matrix proteins; such as Type II collagen and Aggrecan and proteins involved in GAG synthesis (Reviewed in Grad et al., 2011). Biomechanical stimuli have been widely documented to regulate signalling genes involved in bone formation in vivo including constituents of the BMP pathway and *Ihh* (Reviewed in Chen et al., 2010; Nowlan et al., 2008). For mechanical activity to shape the skeleton, alterations to cell behaviour need to occur. A reduction in cell proliferation has been reported in regions of the joint affected morphologically by immobilisation, such as the intercondylar fossa in chick knee joints, mouse mandibular condyles and the joint interzone of *splotch* mice (Jahan et al., 2014; Kahn et al., 2009; Roddy et al., 2011). Cell orientation changes are seen in the jaw cartilages of zebrafish that lack muscle activity (Brunt et al., 2015; Shwartz et al., 2012). Mechanical stimuli are also required for tendon and ligament formation and maturation in species ranging from zebrafish to humans (Chen and Galloway, 2014; Reviewed in Chen and Galloway, 2017). Although cell proliferation and orientation at joints have been shown to be biomechanically controlled, as yet, the signals and pathways that transduce the mechanical stimuli into a cellular response have not been fully elucidated.

Wnts are a family of secreted signalling glycoprotein molecules that play vital roles in development, health and disease (Reviewed in Niehrs, 2012). Classically, Wnt ligands were subdivided into those that activate the canonical beta-catenin pathway or the non-canonical pathways such as Planar Cell polarity (PCP) and calcium-mediated pathways. However, a more recent consensus is that control of the pathway is interlinked and that Wnt ligands can activate multiple pathways depending on tissue type and cellular context (Willert and Nusse, 2012). Many Wnts including Wnt4, Wnt5b and Wnt9a; which typically operate in the canonical pathway, and non-canonical Wnt5a are expressed in developing skeletal elements and are implicated in roles such as regulation of chondrocyte differentiation (Church et al., 2002; Hartmann and Tabin, 2000; Yang et al., 2003) and joint cell identity (Guo et al., 2004; Hartmann and Tabin, 2001). Wnt4, Wnt16, Wnt11 and sFRP2 are all expressed at developing joints (Guo et al., 2004; Ikegawa et al., 2008; Pazin et al., 2012; Rolfe et al., 2014; Witte et al., 2009). Members of the Wnt signalling pathway have also been identified as mechanosensitive. For example, dynamic loading of cultured mesenchymal stem cells affects the regulation of Wnt related genes such as Frizzled-7, Wnt3, Wnt5a and Wnt8 (Arnsdorf et al., 2009; Haudenschild et al., 2009). A decrease in canonical beta-catenin activation was found in ‘muscleless’ *Pax3^sp/sp^Splotch* mouse mutants at the joint (Kahn et al., 2009). A transcriptomic study comparing changes to gene expression in humerus tissue between control and Pax3^sp/sp^ *Splotch* mouse mutants demonstrated that loss of limb muscle led to dysrégulation of 34 members of the Wnt signalling pathway; more genes than any other signalling pathway (Rolfe et al., 2014). These included Wnt ligands, Wnt modulators and Wnt downstream targets. Therefore, Wnt signalling activity in skeletal tissue is mechanosensitive and a candidate pathway to act downstream of mechanical stimuli in skeletogenesis.

Here, we describe cell behaviours that contribute to changes in joint morphology by following live zebrafish joint development under normal or reduced biomechanical conditions. We demonstrate that canonical Wnt activity transduces mechanical signalling to bring about cell behaviours such as proliferation, migration, intercalation and cell morphology changes required to shape the joint. We show that Wnt16 controls cell proliferation and migration specifically in cells at the jaw joint of the lower jaw.

## Results

### Canonical Wnt signalling is active at regions of high strain in the zebrafish lower jaw

Finite Element (FE) models mapping the location of strains acting on the zebrafish lower jaw during mouth opening and closure (Brunt et al., 2015), were utilised to identify signalling activity in areas of high strain. High levels of tensile (Fig. 1A) and compressive (Fig. 1A’) strains from mouth opening muscles are exerted at the anterior of the Meckel’s cartilage (MC) and at the outer region of the jaw joint. During mouth closure muscle activity, high strain is located across the jaw joint interzone (Fig. 1B, B’). The canonical Wnt reporter line *Tg*(*7×TCF.XlaSiam:nlsGFP*), (Moro et al., 2012), reveals cells responding to Wnt are located surrounding the lower jaw at 5 days post fertilisation (dpf) (Fig. 1A”), with a heterogeneous population of GFP-positive (GFP+) cells surrounding the anterior Meckel’s cartilage (MC) and the jaw joints (Fig. 1A”,B”). This localisation of GFP-positive cells shows a strong spatiotemporal correlation with regions of the jaw experiencing high strain.

We therefore studied jaw expression of the Wnt reporter from 3-5 dpf, a time previously identified as critical for joint morphogenesis (Brunt et al., 2015). Using morphology, location and immunohistochemical labelling against ligament and tendon, and chondrocyte markers we identified a heterogeneous population of GFP+ cells (Fig. S1), which included: chondrocytes at the jaw joint (Fig. S1A) and along the palatoquadrate (PQ) (Fig. 1C, c) and ligaments and tendon (Fig. 1C-E, Fig. S1B). Additional GFP+ cells surrounding the jaw joints (Fig. 1C-E, **) were identified as perichondrial joint-associated cells. These Wnt responsive cells at the lower jaw are, therefore, not only located in areas subjected to high levels of tensile and compressive strain but include cell types known to respond to biomechanical stimuli such as chondrocytes and ligaments.

**Fig 1.**
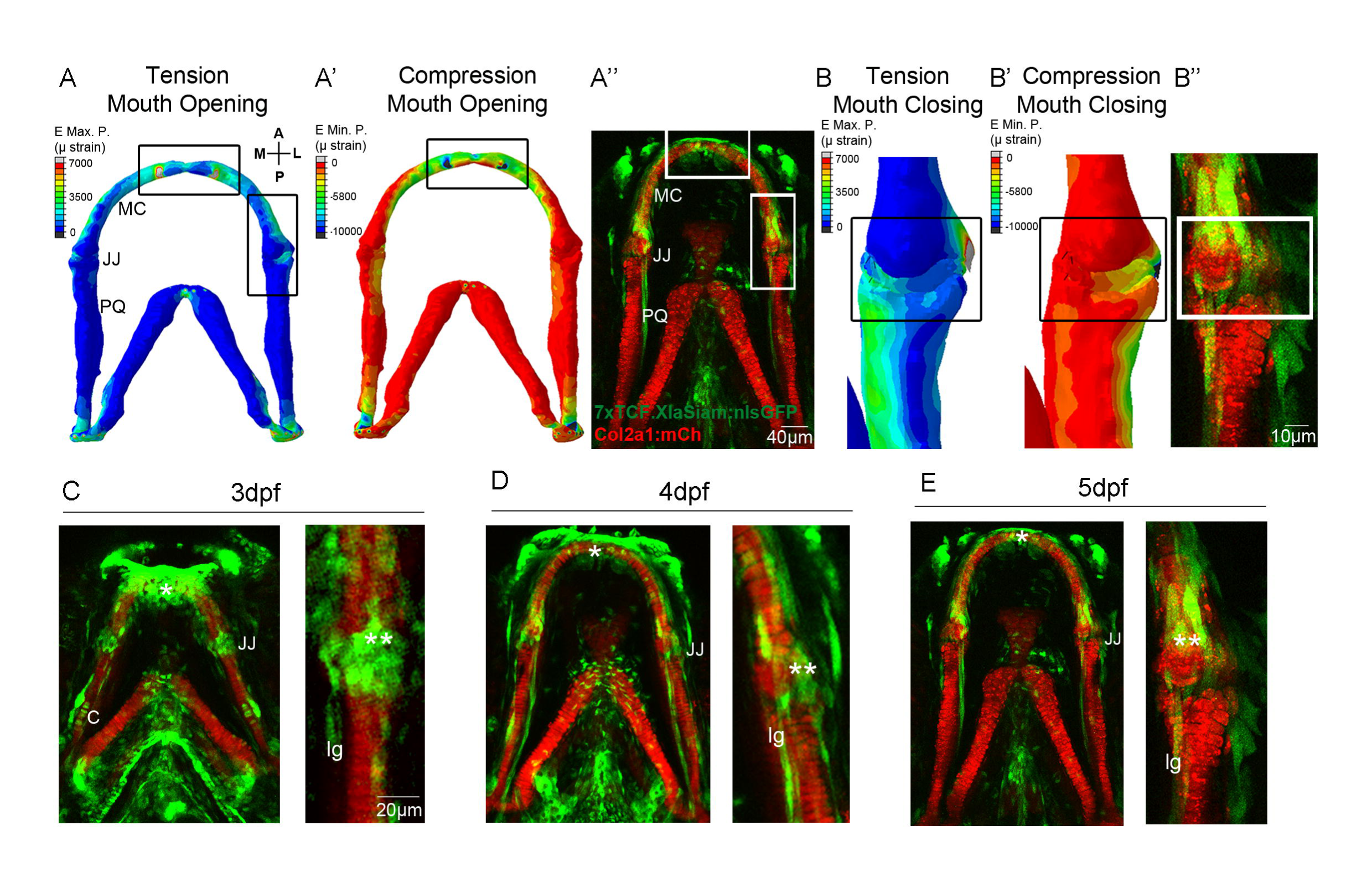
**Patterns of biomechanical strain and location of Wnt responsive cells at the zebrafish lower jaw between 3-5dpf.** (A,A’): Finite Element (FE) model of maximum (E max. P., tension, A) and minimum (E min P., compression, A’) Principal strain on the zebrafish lower jaw during mouth opening at 5dpf.(B,B’): FE model of maximum (E max. P., tension, B) and minimum (E min P., compression, B’) Principal strain on the jaw joint during mouth closure at 5dpf.Colour key represents strain in microstrain (pstrain). (A”,B”,C-E): *Tg*(*7×TCF.XlaSiam:nlsGFP*) and *Tg*(*Col2a1aBAC:mcherry*) transgenic zebrafish lines labelling Wnt responsive cells and cartilage of the lower jaw, respectively at 3 (C), 4 (D) and 5dpf (A”,B”,E). (C-E): left hand panel: lower jaw, right hand panel and (B”): jaw joint. (A,A’) and (B,B’) have been reproduced from the previously published paper under a Creative Commons Licence (Brunt et al., 2015). A= anterior, P= posterior, M=medial, L= lateral, MC= Meckel’s Cartilage, JJ= jaw joint, PQ=palatoquadrate, C= cartilage, lg=ligament, *= anterior MC, **= jaw joint.

### Canonical Wnt signalling at the lower jaw is biomechanically controlled

To test whether canonical Wnt signalling in the jaw is mechanically controlled, zebrafish carrying transgenes for *Col2a1aBAC:mcherry* and *7×TCF.XlaSiam:nlsGFP* were immobilised from 3-5dpf to prevent jaw movement, and *Tg*(*7×TCF.XlaSiam:nlsGFP*) GFP+ signal was quantified by measuring the volume of segmented GFP+ cells within a region of interest at the lower jaw (Fig. 2). A significantly reduced GFP+ signal at the lower jaw at 5dpf was present after a period of immobilisation, most notably at the jaw joint region (Fig. 2A,B), as shown by 3D render of the green channel in the area surrounding the jaw joint and PQ. At 5dpf, the volume of GFP+ signal surrounding the MC joint and PQ (Fig. 2C), and specifically at the jaw joint (Fig. 2D), was significantly reduced in immobilised zebrafish (Fig. 2C’,D’). The total number of GFP+ Wnt responsive cells in the jaw joint region was significantly reduced at 5dpf (Fig. 2E,F) and there were significantly fewer GFP+ ligament/tendon cells at 4 and 5dpf (Fig. 2G). This demonstrates that loss of muscle activity affects canonical Wnt activity at the lower jaw, suggesting that Wnt signalling is biomechanically controlled.

**Fig 2.**
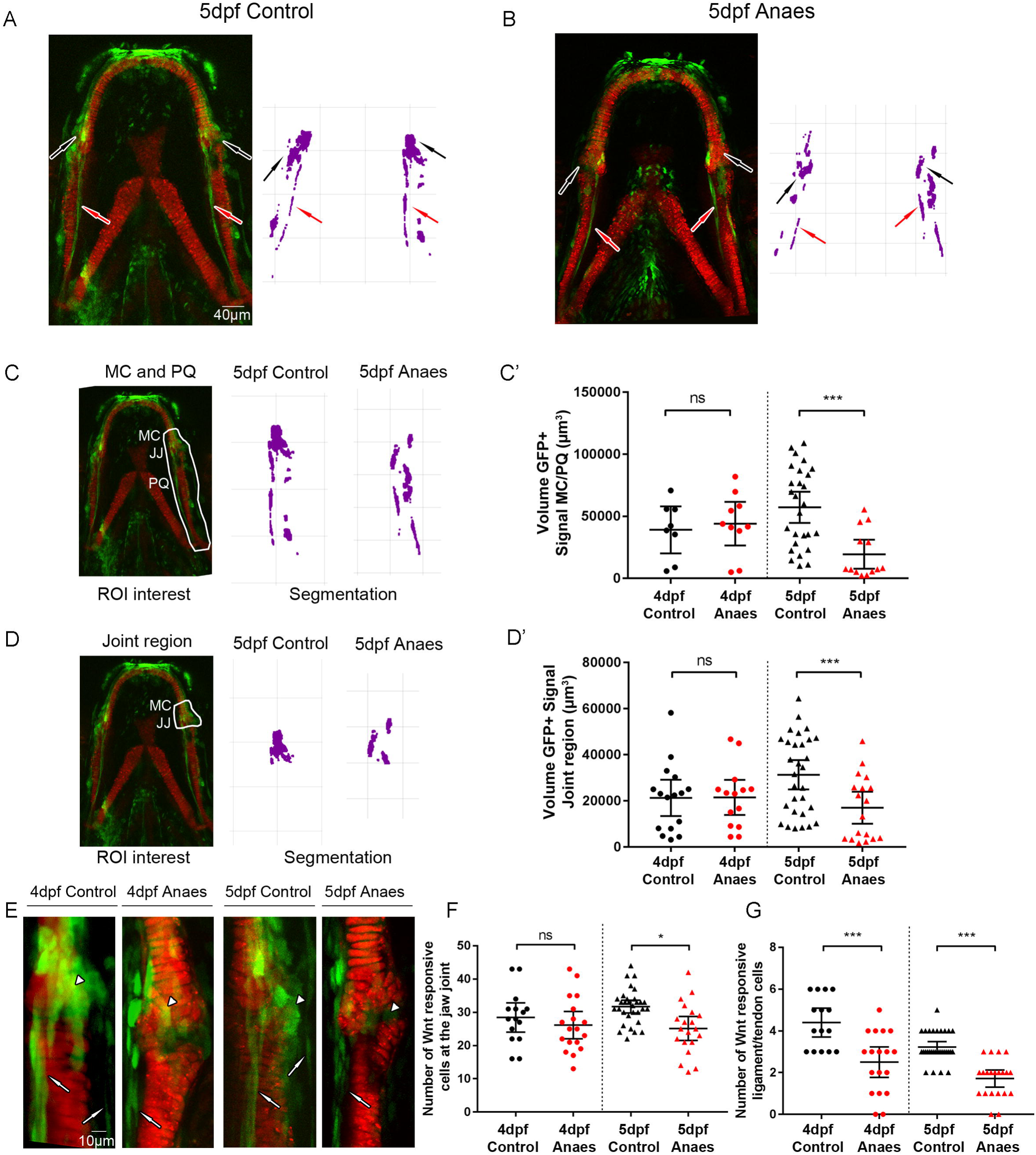
**Immobilisation causes a reduction in canonical Wnt signalling activity at the zebrafish lower jaw.** (A,B): *Tg*(*7×TCF.XlaSiam:nlsGFP*) and *Tg*(*Col2alaBAC:mcherry*) transgenic zebrafish lines were used to visualise Wnt responsive cells and chondrocytes, respectively in 5dpf control (A) and 5dpf 3-5dpf immobilised (B) zebrafish. Left panel= merge of *Tg*(*7×TCF.XlaSiam:nlsGFP*) and *Tg*(*Col2alaBAC:mcherry*). Right panel= Segmentation of GFP signal. Black arrows= Cells surrounding jaw joint, red arrows= ligaments and tendons. (C): Left panel: volume analysis of *Tg*(*7×TCF.XlaSiam:nlsGFP*) GFP-positive (GFP+) signal at the region of interest (ROI): from 6 intercalating cells above the Meckel’s Cartilage (MC) jaw joint and along the full extent of the palatoquadrate (PQ) (white line). Right panel: Segmentation of GFP+ signal volume from region of interest in 5dpf control and anesthetised zebrafish. (C’): Volume (μm^3^) of GFP+ signal at the MC joint and along the PQ in 4 and 5dpf control and anaesthetised zebrafish. (n=8, 10, 27, 13 joints). (D): Left panel: volume analysis of *Tg*(*7×TCF.XlaSiam:nlsGFP*) GFP-positive (GFP+) signal at the ROI from 6 intercalating cells above the Meckel’s Cartilage (MC) jaw joint to the interzone (white line). Right panel: Segmentation of GFP+ signal volume from region of interest in 5dof control and anaesthetised zebrafish. (D’): Volume (μm^3^) of GFP+ signal at the MC joint in 4 and 5dpf control and anaesthetised zebrafish. (n=16, 14, 30, 18 joints). (E): *Tg*(*7×TCF.XlaSiam:nlsGFP*) and *Tg*(*Col2alaBAC:mcherry*) transgenic zebrafish mark Wnt responsive cells and cartilage of the lower jaw at the jaw joint in 4 and 5dpf control and anaesthetised zebrafish. White arrowheads indicate joint-associated GFP+ cells. White arrows indicate ligament and tendon GFP+ cells. (F): Number of GFP+ cells in 4 and 5dpf control and anaesthetised zebrafish in a 50x80pm area surrounding the jaw joint. (n=15, 18, 31,13 joints). (G): Number of ligament and tendon GFP+ cells in 4 and 5dpf control and anaesthetised zebrafish at the jaw joint. (n=15,18, 31, 13 joints). Kruskal-Wallis tests were performed for statistical analysis in (C’,D’,G) and one-way ANOVA in (F). ns= not significant, *=p≤0.05, **=p≤0.01, ***=p≤0.001. Bars on graph represent mean and 95% confidence interval (Cl).

### Blocking canonical Wnt signalling leads to altered jaw joint morphology independent of muscle activity

We have previously shown that immobilising the jaw leads to abnormal joint formation (Brunt et al., 2015; Brunt et al., 2016b). To test whether changes to Wnt activity affect jaw joint morphology, independent of movement, *Tg*(*Col2a1aBAC:mcherry*) zebrafish were exposed to the Wnt antagonist, IWR-1, from 3-5dpf. The addition of IWR-1 had no significant effect on the frequency of mouth movements compared to control (Fig. S2), i.e. jaw muscle activity was normal. However, IWR-1 treatment affects the functional morphology of the 5dpf jaw joint, such that the medial region of the MC overlapped the PQ element, impeding smooth movement (Fig. 3A,A’). IWR-1 treatment caused the lateral interzone region to be significantly larger than control and the medial interzone region to be significantly reduced due to overlapping elements (Fig. 3B-B’). There was no effect on the total length of the jaw (Fig. S3A) or MC (Fig. S3B), suggesting that normal growth was not inhibited. However, the proportion of chondrocytes in the MC that were fully intercalated was significantly reduced (Fig. S3C), concurrent with a significant increase in the proportion of rounded chondrocytes at the 5dpf jaw joint (Fig. S3D). This failure of intercalation and reduced cell maturation at the joint phenocopies what is previously seen in immobilised *nic* b107 mutants and anaesthetised zebrafish (Brunt et al., 2016b; Shwartz et al., 2012). Therefore, IWR-1 treatment, independent of muscle activity and joint movement, recapitulates cell behaviours seen after immobility.

**Fig 3.**
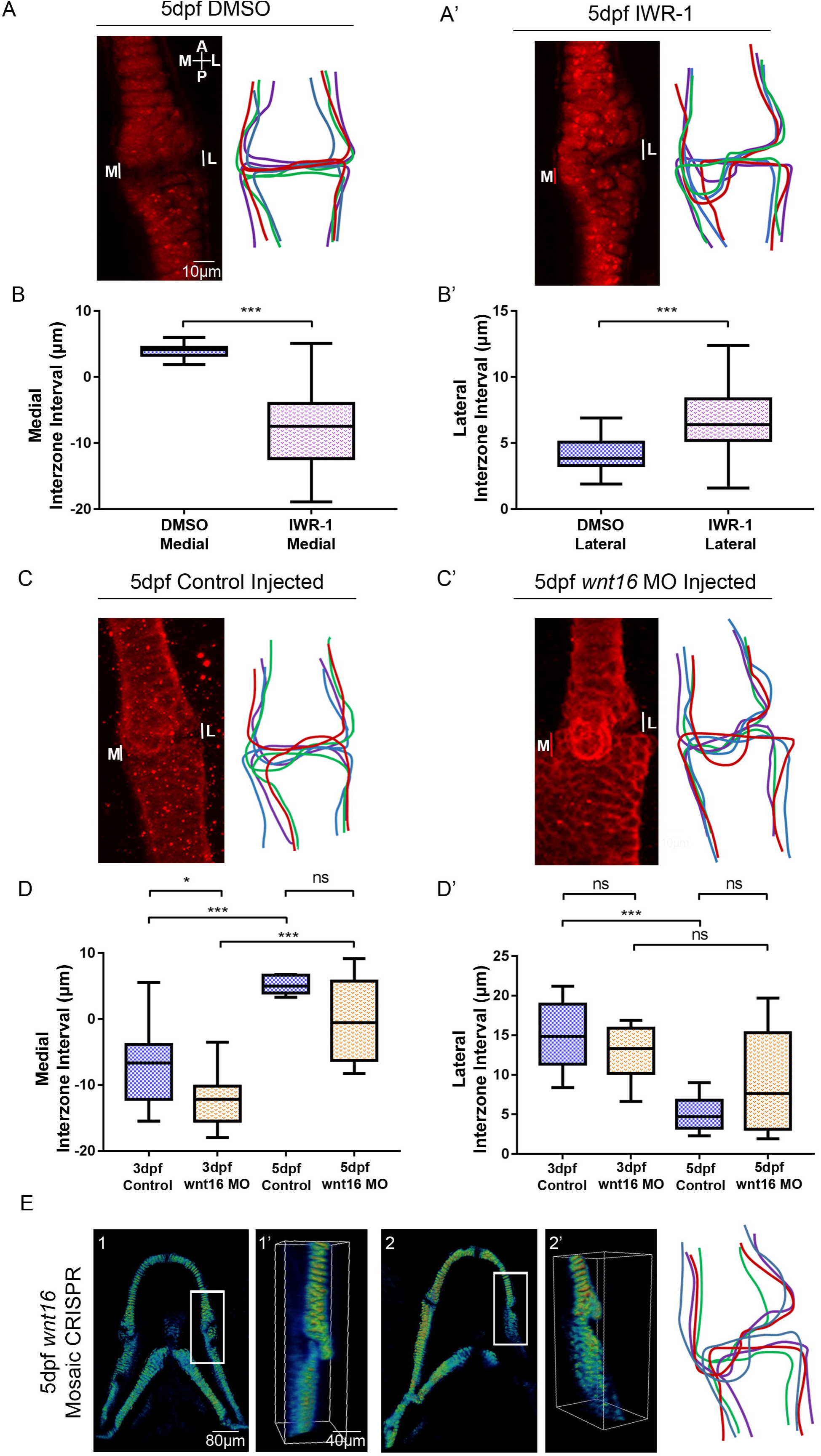
**Manipulation of Wnt affects zebrafish jaw joint morphology.** (A-A’): 5dpf DMSO control (A) and 20pM IWR-1 treated (A’) zebrafish jaw joint morphology. Left panels: *Tg*(*Col2alaBAC:mcherry*) transgenic zebrafish line marks cartilage of the jaw joint. Right panels: Outlines of 4 representative jaw joints. A= Anterior, P= Posterior, M= Medial, L= Lateral. White lines= Interzone Interval measurements between MC and PQ, red line= Overlapping Interval between MC and PQ. (B-B’): Interzone intervals (μm) between the MC and PQ on the medial (B) and lateral (B’) regions of the jaw joint in 5dpf DMSO and IWR-1 treated zebrafish. Negative values represent an overlap of MC/PQ elements. (n=42, 45 joints). Two-tailed student t-tests were performed for (B,B’). (C-C’): 5dpf control injected (C) and Wnt16 morpholino (MO) injected (C’) zebrafish jaw joint morphology. Left panels: Immunohistochemical stain of the jaw joint region. Right panels: Outlines of 4 representative jaw joints. (D-D’): Interzone intervals (μm) between the MC and PQon the medial (D) and lateral (D’) regions of the jaw joint in 5dpf control injected and Wnt16 MO injected zebrafish. Negative values represent an overlap of MC/PQ elements. (n=8,11, 6, 8 joints). One-way ANOVAS were performed (D,D’). ns= not significant, *=p≤0.05, **=p≤0.01, ***=p≤0.001. Bars on graph represent mean and 95%CI. (E): 3D volume rendering of 5dpf injected CRISPR/Cas9 mosaic *Wnt16* knockout TgBAC(*col2al:mCherry*) larvae (1-2), (n= 12 animals). Image was zoomed in and rotated to best show the jaw joint (l’−2’). Right panel: Outlines of 4 representative jaw joints.

### Knockdown and mosaic knockout of Wnt16 leads to altered jaw joint morphology

Application of a Wnt antagonist (IWR-1) shows that a reduction in canonical Wnt activity leads to abnormal jaw joint morphogenesis. We took a candidate approach to identify Wnt pathway members that could transduce the mechanical signal into altered cell behaviour. Wnt16 has been previously reported to be expressed in mouse limb joints (Witte et al., 2009), differentially regulated in mice lacking limb muscle (Rolfe et al., 2014) and Wnt16 overexpression in mouse joint synovium has been shown to activate canonical Wnt signalling in joint cartilage (van den Bosch et al., 2015). In zebrafish, Wnt16 is required to control notch signalling for haematopoietic stem cell specification (Clements et al., 2011). We used the previously described Wnt16 MO, (Clements et al., 2011), to determine the effect of reduced Wnt16 on jaw and joint morphology. Wnt16 knockdown had no effect on the gross morphology of the zebrafish larvae (Fig. S4B) or on frequency of jaw movement (data not shown). Wnt16 knockdown led to reduced levels of *lef1* mRNA in jaw cartilage elements such as the branchial arches (Fig. S5A,A’), whilst leaving other expression domains - such as the brain - intact (Fig. S5A,A’). Wnt16 morphants showed a significant reduction in *Tg*(*7xTCF.XlaSiam:nlsGFP*) GFP+ signal volume in the jaw compared to control (Fig. S5B-E), demonstrating that Wnt16 activates canonical Wnt signalling in the lower jaw independent of jaw movement.

Wnt16 morphants show altered jaw joint morphology with an overlapping MC element (Fig. 3C-C’). At 3dpf, Wnt16 morphants have a reduced interzone space on the medial side of the joint due to the overlap of the elements on the medial side (Fig. 3D-D’). Interzone interval measurements were, however, not significantly different to control at 5dpf. This shows Wnt16 knockdown affects functional jaw joint morphology, but is less severe than IWR-1 treated fish at 5dpf. Wnt16 MO injection did not significantly affect the total jaw length or MC length at 5dpf, showing that overall jaw growth was unaffected and Wnt16 knockdown very specifically affects only the joint region of the cartilage element (Fig. S3A,B), with no other discernible off-target effects. Mosaic Wnt16 knockout also leads to abnormal jaw joint morphology with an overlapping medial joint region, but also does not affect normal jaw growth and development (Fig. 3E). Unlike IWR-1 treatment, Wnt16 knockdown and mosaic knockout does not significantly affect cell intercalation (Fig. 3E, Fig. S3C,D). Therefore, Wnt16 is important for joint morphology, but does not affect cell intercalation.

### Cell proliferation, migration and changes to cell morphology contribute to jaw joint morphogenesis

In order to understand the cell behaviour changes that shape the joint, we tracked cells at the joint in individual larvae. As continuous time lapse imaging to follow the process of joint morphogenesis would require long-term immobilisation; which would in turn lead to abnormal morphogenesis, we used zebrafish carrying both *Tg*(*Sox10:GAL4-VP16*) and *Tg*(*UAS:Kaede*) transgenes to track populations of kaede-expressing joint cells from 3 to 5dpf. A small batch of 10-12 cells at the medial side of the joint were photoconverted at 3dpf to irreversibly switch the labelling from green to red, making it possible to follow the cells over time. Medially located cells close to the retroarticular process (RAP) were chosen as the medial region of the joint is most affected by immobilisation and Wnt abrogation (Fig. 3, (Brunt et al., 2015)). Red photoconverted cells in the medial joint at 3dpf spread along the anterior-posterior axis of the jaw joint by 5dpf, contributing to the change in joint shape (Fig. 4A,C). Some cells within this group remain part of the MC; however other cells migrate to the PQ. Between 3 and 5dpf there is a 97.5% mean increase in the area occupied by red cells (Fig. 4E). Cell counts reveal that this area increase, in part, is due to an increase in cell number from 3-5dpf (Fig. 4F), showing that proliferation contributes to changes in joint shape. BrdU pulse chase experiments show that proliferation events mainly occur between 4 and 5dpf (Fig. S6A-A”,D). From 3 to 5dpf, cell morphology changes are also observed, with elongated perichondrial cells migrating from the original pool of round photoconverted cells to form the perichondrium (Fig. 4A,C). These data demonstrate that cells at the joint are highly dynamic, with migration, proliferation and changes to cell type and morphology all contributing to normal joint morphogenesis.

We then tracked joint cells in immobilised larvae to investigate whether cell behaviour is altered in these larvae. In immobilised larvae, red photoconverted cells remained largely static between 3 and 5dpf (Fig. 4B) and the percentage increase in the area occupied by red cells was significantly reduced compared to control (Fig. 4E). The percentage increase in the number of cells inheriting red kaede at the joint between 3 and 5 dpf was also significantly reduced (Fig. 4F). Therefore, mechanical stimuli are required to trigger normal cell behaviours such as proliferation and migration in order to correctly shape the joint.

**Fig 4.**
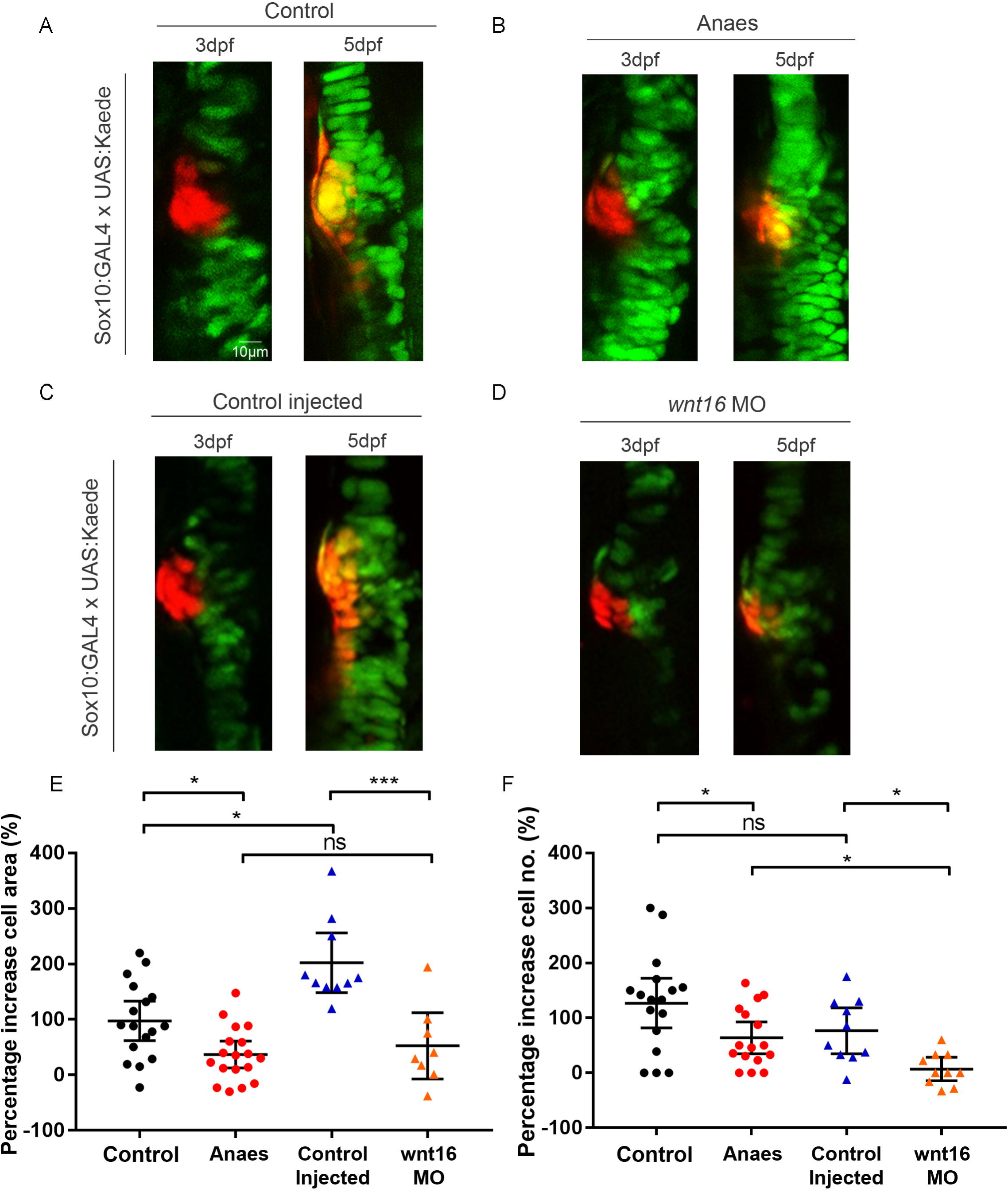
**Immobilisation and Wnt16 knockdown affects cell proliferation and migration at the medial region of the jaw joint between 3-5dpf.** (A-D): *Tg*(*Sox10:GAL4-VP16*) and *Tg*(*UAS:Kaede*) transgenic line drives expression of kaede protein (green) in cartilage of control (A), anaesthetised (B), control injected (C) and Wnt16 morpholino (MO) injected (D) zebrafish. At the jaw joint, medially located kaede expressing cells are photoconverted to red kaede at 3dpf (left panels). Right panels show jaw joints from the same larva reimaged at 5dpf. Photoconverted cells appear red/orange due to presence of photoconverted red kaede and expression of newly made green kaede protein under control of sox10 promoter. (E): Percentage increase in total area of cells expressing photoconverted red kaede between 3 and 5dpf in control, anaesthetised, control injected and Wnt16MO injected zebrafish jaw joints. (n=17, 18,10, 8 joints). (F): Percentage increase in number of cells expressing photoconverted red kaede between 3 and 5dpf in control, anaesthetised, control injected and Wnt16MO injected zebrafish jaw joints. (n=17, 16,10,10 joints). Kruskal-Wallis tests were performed for (E,F). ns= not significant, *=p≤0.05, **=p≤0.01, ***=p≤0.001. Bars on graph represent mean and 95%CI.

### Wnt16 controls cell proliferation and migration at the jaw joint

To investigate whether Wnt16 plays a role in controlling cell behaviours at the joint, a group of 10-12 red photoconverted cells per fish were tracked in Wnt16 morphants and Wnt16 mosaic CRISPR knockouts. From 3 to 5dpf, the spread of red photoconverted cells observed in control injected larvae did not take place in Wnt16 morphants and Wnt16 mosaic CRISPR knockouts (CRISPants) (Fig. 4C,D, Fig. S7A,B). The percentage increase in red cell area was significantly reduced in morphants and CRISPants compared to control larvae (Fig. 4D,E, Fig. S7C). The percentage increase in cell number was significantly reduced in morphants compared to control (Fig. 4F) and the number of BrdU positive cells at the joint was also significantly fewer (Fig. S6C,D). This shows that Wnt16 controls cell behaviours including proliferation and migration during joint morphogenesis. Interestingly, the effect of Wnt16 knockdown on chondrocyte migration and proliferation is highly joint specific, as there was no significant change in cell behaviour in the more mature intercalated region (Fig. S7B’,C’,E, Fig. S8); further confirming specificity of the MO and CRISPR.

Next, we used zebrafish with the *ubi:Zebrabow* transgene under the control of *Sox10:cre* to track individual cells, in order to unpick individual cell behaviours taking place during joint morphogenesis. At 3dpf, the retroarticular process (RAP) contains cells with different colour profiles, which could be tracked (Fig. 5A-A”). In control zebrafish, cells that had undergone proliferation between 3-5dpf were observed at the joint (Fig. 5B, duplicated white and green asterisks). Cell morphology changes were also observed (Fig. 5B’, yellow and blue outlined cells).

However, in IWR-1 treated and Wnt16 morphant larvae, cell proliferation was not observed in the RAP between 3-5dpf (Fig. 5C-D). In IWR-1 treated zebrafish the cells of the RAP are less plastic, with minimal changes to cell morphology between 3-5dpf (Fig. 5C’). In Wnt16 morphants, cell morphology changes were not affected, as the cells of the RAP became enlarged or changed shape (Fig. 5D’, e.g. lower blue cell). Both cellular processes are affected by IWR-1 application and proliferation is affected by Wnt16MO knockdown. This shows that Wnt16 controls cell proliferation and migration at the joint, but suggests cell morphology changes may be controlled by other members of the Wnt pathway.

**Fig 5.**
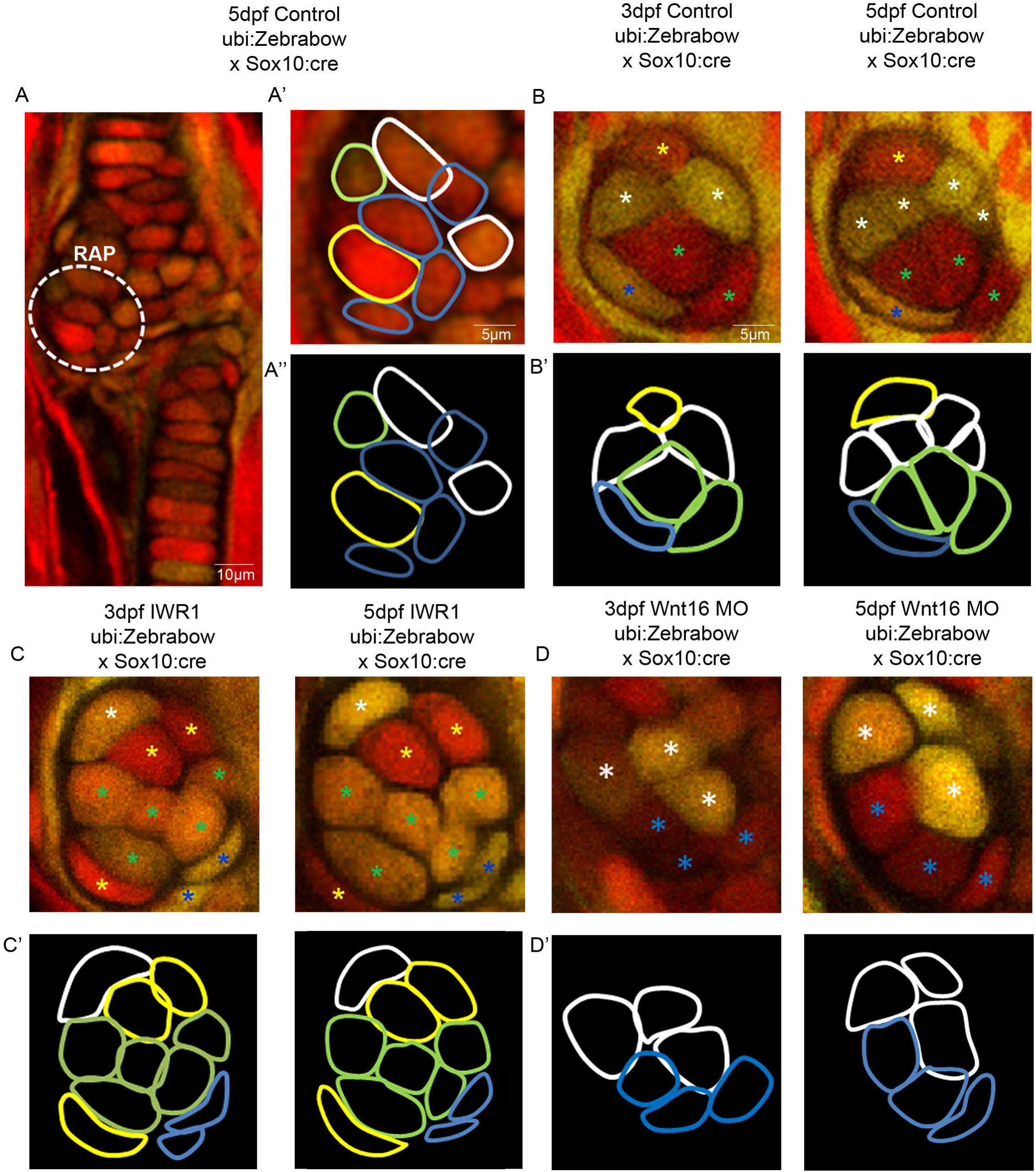
**Wnt manipulation affects cell proliferation and cell morphology at the jaw joint revealed using Zebrabow transgenic line.** (A-A”): *Tg*(*ubi:Zebrabow*) and *Tg*( *Sox10:cre*) transgenic lines generate multiple colours of fluorescence in zebrafish cartilage including at the region of interest at the Retroarticular process (RAP), (white dotted line). Cell outlines were created at the RAP (A’: RAP cell outlines overlay with confocal image, A”: cell outlines). A=anterior, P=posterior, M=medial, L=lateral. (B, C, D): 3 and 5dpf *Tg*(*ubi:Zebrabow*) and *Tg*(*Sox10:cre* control (B), IWR-1 treated (C) and Wnt16 MO injected zebrafish (D). The Retroarticular process (RAP) of the MC jaw joint is shown. Asterisks of different colours mark cells at 3 and 5dpf (indicating re-identification and cell division events). (B’, C’, D’): Outlines of individual cells in the RAP of the MC jaw joint, identified in Control (B’), IWR-1 treated (C’) and Wnt16 MO injected (D’) *Tg*(*ubi:Zebrabow*) and *Tg*( *SoxlOxre*) transgenic zebrafish. Outline colour of individual cells in (B’, C’, D’) matches asterix colour in (B, C, D).

## Discussion

Mechanical input has previously been shown to affect joint morphogenesis in a number of species ranging from mouse to fish (Brunt et al., 2015; Brunt et al., 2016b; Kahn et al., 2009; Nowlan et al., 2010b; Rolfe et al., 2013). However, which downstream signalling pathways drive the cell behaviours that shape the joint in response to these forces is less well characterised. By tracking cell behaviour dynamically in the joint for the first time in larvae subjected to mechanical, genetic and pharmacological perturbations, we show that joint morphology is shaped through a combination of cell morphology changes, migration and proliferation. Cells in the medial region of joint; most affected in mechanical loss models, normally spread and migrate anterior and posterior to their original location to remain part of the Meckel’s cartilage or become part of the palatoquadrate. Cell proliferation in the jaw joint mainly occurs from 4 to 5dpf and cell morphology changes also contribute to the overall shape of the joint. We also show that interzone cells can give rise to mature chondrocytes or form the perichondrium. This is the first study to describe the dynamic cell behaviours occurring in joints in individually tracked animals and therefore gives a dynamic insight into morphogenesis of the joint *in vivo.* We show that removal of muscle force leads to reduced proliferation in the zebrafish joint, analogous to the situation in chicks and mice (Jahan et al., 2014; Kahn et al., 2009; Roddy et al., 2011). Our work also builds on previous work showing the relevance of the zebrafish as a model for synovial joint development (Askary et al., 2016).

Here, we demonstrate that canonical Wnt signalling, and Wnt16 act downstream of muscle activity to transduce the mechanical signals into the cell behavioural changes, such as proliferation and migration, that shape the joint. It has been previously shown in mesenchymal stem cell *in vitro* preparations that mechanical strain can activate Wnt signalling (Arnsdorf et al., 2009; Haudenschild et al., 2009; Rolfe et al., 2014) and transcriptomic studies in muscle-less mice have demonstrated changes to expression of Wnt pathway members (Rolfe et al., 2014). We show that canonical Wnt signalling is activated in cells associated with the zebrafish jaw joint, which are located in regions under high levels of strain. We demonstrate that canonical Wnt signalling in the jaw joint, and in ligaments is mechanosensitive, with significant reductions to the number Wnt GFP+ positive cells at the joint and in associated connective tissues when force is lost. Previous work in zebrafish has shown that craniofacial muscle is not required for induction of expression of early markers of tendon and ligament, but that muscle attachment is required for maintenance of expression at 72hpf (Chen and Galloway, 2014). Immobilisation in our study starts somewhat later (from 72hpf) but we can still identify changes in the ligaments and a reduction of Wnt signal activity at 4 and 5dpf. This strongly suggests that I Wnt signalling plays a mechanosensitive role in later tendon and ligament maturation. Wnt/β-catenin has been linked to a mechanosensitive role in controlling expression of osteogenic genes in cells derived from human periodontal ligaments (Chen et al., 2017; Zhang et al., 2016), and our work would suggest that Wnt is likely to play a role in maturation of other craniofacial ligaments. We observe that canonical Wnt signalling manipulation causes abnormalities in joint morphology. This occurs even in conditions where muscle activity is still present, demonstrating that that Wnt signalling acts downstream of muscle activity to cause changes in joint shape.

We show that Wnt16 is important for accurate shaping of the joint by controlling cell proliferation and migration events at the joint. Unlike reduction in broad canonical Wnt signalling, abrogation of Wnt16 had no effect on cell behaviours such as proliferation, migration and intercalation of maturing chondrocytes anterior to the jaw joint, acting in a highly joint specific fashion. Therefore, other Wnt ligands are likely to be responsible for chondrocyte intercalation in the Meckel’s cartilage, as is the case for chondrocyte intercalation in chick growth plates (Li and Dudley, 2009; Rochard et al., 2016) and during zebrafish palate morphogenesis (Dougherty et al., 2013; Kamel et al., 2013; Rochard et al., 2016). Wnt16 has been previously implicated in joint formation, bone homeostasis and remodelling (Gori et al., 2015; Guo et al., 2004; Kobayashi et al., 2016) and is expressed at the developing joint in mouse models (Guo et al., 2004; Witte et al., 2009). It has been shown to be required for proliferation, differentiation and specification in other cell types such as haematopoietic stem cells, osteoclasts, osteoblasts and keratinocytes (Clements et al., 2011; Kobayashi et al., 2015; Ozeki et al., 2016; Teh et al., 2007). Wnt16 is upregulated following mechanical injury in *ex vivo* human cartilage (Dell’Accio et al., 2008), and following mechanical loading of the tibia in mice (Wergedal et al., 2015). Expression levels of Wnt16 were upregulated in ‘muscleless’ *Splotch* mice compared to control (Dell’Accio et al., 2008; Rolfe et al., 2014; Wergedal et al., 2015). We show that Wnt16 controls proliferation and migration of a small number of cells in the joint, which are critical for normal joint morphology to be generated.

The role of Wnts; in particular Wnt16, by controlling joint morphogenesis during development may have a longer term impact on joint health. The formation of abnormal joint morphology during development is a critical risk factor in onset of osteoarthritis (Baker-LePain and Lane, 2010). Wnt-related genes such as Wnt antagonist FRZB are implicated in accurate joint shaping (Baker-Lepain et al., 2012). Wnt16 has been linked with the relationship between hip geometry and risk of osteoarthritis onset (Garcia-Ibarbia et al., 2013). Wnt16 is upregulated in joints with moderate to severe OA along with increased nuclear beta-catenin expression (Dell’Accio et al., 2008). Upregulation is also documented after mechanical injury (Dell’Accio et al., 2008). Our study builds on these findings to suggest that the relationship found between OA risk, joint shape and Wnt16 may stem from its role in activating early joint cell behaviours that affect the functional joint shape.

## Materials and Methods

### Zebrafish husbandry and transgenic lines

Zebrafish were maintained as previously described (Westerfield, 2000). Experiments were approved by the local ethics committee and granted a UK Home Office project licence. Transgenic lines *Tg*(*7×TCF.XlaSiam:nlsGFP*) (Moro et al., 2012), *Tg*(*Col2a1aBAC:mcherry*) (Hammond and Schulte-Merker, 2009), *Tg*(*Sox10:GAL4-VP16*) (*Lee et al., 2013*), *Tg*(*UAS:Kaede*) (Hatta et al., 2006), *Tg*(*ubi:Zebrabow*) (Pan et al., 2013) and *Tg*(*−4.7Sox10:cre*) (Rodrigues et al., 2012) have been previously described. Larvae from the same lay were randomly assigned to different treatment groups.

### Pharmacological treatment

Fish were anaesthetised between 3 and 5dpf with 0.1mg/ml MS222 (Tricaine methanesulfonate) (Sigma) diluted in Danieau solution. MS222 and Danieau solution was refreshed twice daily. 20μΜ IWR-1 (Sigma), was diluted in Danieau solution and replaced daily.

### Finite Element (FE) models

Meshes for 5dpf FE models have been previously published (Brunt et al., 2015; Brunt et al., 2016a). Loads for jaw opening (Protractor Hyoideus and Intermandibularis Anterior muscles) and jaw closure (Adductor Mandibulae muscles) were applied to predict tensile and compressive strains. FE results are displayed as colour contour plots of maximum and minimum strain.

### Wnt responsive cell counts and area

Image stacks of 4 and 5dpf *Tg*(*7×TCF.XlaSiam:nlsGFP*) *xTg*(*Col2a1aBAC:mcherry*) transgenic jaws were imported into Fiji (Schindelin et al., 2012). Wnt responsive cells of ligament and tendon morphology along the palatoquadrate (PQ) element were counted. All Wnt responsive cells surrounding the jaw joint within a 50×80 μm area were counted.

A custom script was written in MATLAB (version 2015a; Mathworks, Inc.) so that a selected area of *Tg*(*7xTCF.XlaSiam:nlsGFP*) GFP positive signal could be determined. Areas of interest included: 1. the area of the Meckel’s Cartilage (MC) (from 6 intercalating cells above the MC joint) plus the PQ, 2. the joint region (from 6 intercalating cells above the MC joint, to the MC interzone). Coarse regions of interest were initially manually identified from maximum intensity projections of the image stack and subsequently segmented in 3 dimensions (3D) based on the MATLAB implementation of Otsu’s threshold (Otsu, 1979). All voxels within a user-selected region of interest with intensity values above the threshold were classified as a single object. An alpha shape was calculated for each segmented object (Edelsbrunner et al., 1983) using MATLAB’s automatically determined surface radius. Volumes for each object were measured using the method provided by the MATLAB alpha shape class.

### Wholemount Immunohistochemistry

Wholemount immunohistochemistry was carried out as previously described (Hammond and Schulte-Merker, 2009). Larvae to be stained for BrdU were treated with 2N HCl for 1hr at 37°C. The following primary antibodies previously used in zebrafish (Table S1.) were used: (Chicken anti-GFP, ab13970, Abcam, 1:500 dilution; rabbit anti-tenascin C, USBI142433, US biological, 1:300 dilution; mouse anti-BrdU, B8434, Sigma, 1:100 dilution; rabbit anti-collagen II, ab34712, abcam, 1:200 dilution; mouse anti-collagen II (II-II6B3), AB528165, Developmental Studies Hybridoma Bank, 1:200 dilution). Secondary antibodies: (Dylight 550 goat anti-mouse IgG, 84540; Dylight 488 goat antimouse IgG, 35502; Dylight 550 goat anti-rabbit IgG, 84541; Dylight 488 goat anti-chicken, IgY, SA5- 10070; ThermoFisher Scientific, 1:500 dilution).

### Joint outline and interzone interval analysis

Tiff images of *Tg*(*Col2a1aBAC:mcherry*) transgenic labelled joints were imported into Powerpoint. The draw tool was used to draw around 4 representative joints for each condition and overlaid for analysis.

The interval between MC and PQ cartilage elements on the medial and lateral side of the jaw joint were measured from tiff images in LAS AF Lite software. Negative values correspond to instances of overlapping cartilage elements.

### Kaede protein photoconversion

Double transgenic *Tg*(*Sox10:GAL4-VP16*) *× Tg*(*UAS:Kaede*) zebrafish larvae at 3dpf were mounted ventrally on coverslips in 0.3% agarose under MS222 anaesthetic. The FRAP wizard setting on Leica LAS software was used to photoconvert kaede expressing cells of interest from green to red fluorescence on a Leica SP5 or SP8. Briefly, a region of interest (ROI) was drawn using the selection tools, on the medial Meckel’s cartilage joint or the intercalating cell region of the MC. The 405nm was used to photoconvert cells in the ROI at 8% laser power for 10 seconds. Following photoconversion, larvae were removed from agarose and flushed with Danieau solution until resumption of movement. Each larva was kept separately for individual identification. Larvae were left to develop normally or anaesthetised with MS222 and reimaged at 5dpf. Daughter cells inherit irreversibly photoconverted red kaede protein after cell division (Mutoh et al., 2006).

### Photoconverted cell number and area change

Image stacks containing the red channel were imported into Fiji software (Schindelin et al., 2012) and red cell numbers were counted at 3 and 5 dpf, and percentage increase in cell number calculated.

The image stacks were saved as a tiff file. A Fiji plugin designed to segment a thresholded level of red cells was used to calculate the combined area of red cells; these areas were compared from the individual larvae from 3 to 5 dpf. The percentage increase in cell area was calculated.

### Jaw and element length and ratio of cell type in MC element

Confocal images of jaw joints labelled with Tg(Col2a1aBAC:mcherry) were loaded into Fiji (Schindelin et al., 2012). The length (μm) of the jaw was measured from anterior MC to posterior palatoquadrate using the line tool. The length of the MC element was measured using the freehand line tool from anterior MC to the MC jaw joint. The proportion of the length (μm) of the MC comprising rounded cells or intercalating cells was measured in Fiji using the freehand line tool. The ratio of the length of the MC occupied by varying cell types compared to the full MC length was then calculated.

### BrdU

Larvae were treated with 3mM BrdU (Sigma) diluted in Danieau solution from 3-4dpf or 4-5dpf. After treatment, larvae were washed 4×5min in Danieau solution then fixed with 4% PFA overnight at 4°C. Larvae were immunohistochemically stained for BrdU (as detailed above).

### Zebra bow

Double transgenic (*Tg*(*Ubi:Zebrabow)*) and *Tg*(*Sox10:Cre*) zebrafish larvae express a variety of different fluorescent protein combinations in the cells of the developing cartilage. This allows individual cells to be tracked as they migrate or divide. 3dpf *Tg*(*Sox10:Cre*) × *Tg*(*Ubi:Zebrabow*) double transgenic zebrafish were mounted in 0.3% agarose on coverslips in dishes and covered with Danieau solution containing MS222. The larvae were imaged on a Leica Multiphoton microscope using a 25x water dipping lens. Three fluorescence channels were collected individually (YFP, RFP and GFP). Larvae were returned to Danieau solution in individual dishes and either left to develop normally or anaesthetised with MS222 until 5dpf, then reimaged.

### Wnt16 Morpholino knockdown

A Wnt16 splice-blocking morpholino (MO) (Gene-Tools), AGGTTAGTTCTGTCACCCACCTGTC was used to knockdown Wnt16 protein as previously described (Clements et al., 2011). 5ng of Wnt16 or control morpholino was injected with rhodamine dextran and 0.2M potassium chloride into 1 cell stage *Tg*(*SoxlO:GAL4-VP16*) *× Tg*(*UAS:Kaede*) embryos using a picospritzer III (Parker) microinjector.

### RNA extraction and making Wnt16 cDNA

Failure of splicing after Wnt16 MO injection was confirmed by PCR (Fig. S4A). Total RNA was extracted from pooled and homogenised 3dpf Wnt16 MO injected and control non-injected larvae using a Nucleospin RNA II kit, (Macherey-Nagel). cDNA was produced from 1μg RNA via reverse transcription using M-MLV reverse transcriptase (Promega). cDNA was amplified by PCR using Wnt16 primers (forward: ACTAAAGAGACAGCTTCATCC, reverse: AACTCATCTTTGGTGATAGGC, (Eurofins Genomics)) (Clements et al., 2011) and Taq polymerase (Roche). PCR conditions are previously described (Clements et al., 2011).

### Wnt16 CRISPR mosaic knockout

CRISPR target sequences were selected using CRISPscan track from UCSC Browser (danRerlO), (Moreno-Mateos et al., 2015), and based on high scores and proximity to Wnt16 first exon (Fig. S9A) two sequences targeting exon 2 were selected: guide1- GGAGGAGTGCCCGAGAAGTT (score 73-chr4:10708367-10708389) and guide2- GGTGGAACTGCTCGACCCGA (score 64-chr4:10708367-10708389). gRNA antisense oligonucleotide sequences (5’−3’) were designed as follows: AAAGCACCGACTCGGTGCCACTTTTTCAAGTTGATAACGGACTAGCCTTATTTTAACTTGCTATTTCTAGCTCTAAAAC - N20 -**CTATAGTGAGTCGTATTA**CGC, with the T7 promoter shown in bold and N20 indicating the reverse complement of targeting sequence, as described previously (Hruscha et al., 2013). *In vitro* transcription was carried out annealing gRNA antisense oligonucleotide to T7 primer (TAATACGACTCACTATAG; 5 minutes at 95°C, cooled at room temperature) followed by transcription using the Ambion MEGAshortscript-T7 kit. Injection mix was prepared to a final concentration of 200 ng/uL of gRNA and 600 ng/uL of GeneArt Platinum Cas9 nuclease (Invitrogen) and incubated for 10 min at RT. 1pL of the solution was injected into the cell of eggs at 1 cell stage (Fig. S9B). To check gRNA efficiency, DNA was extracted from individual larvae at 48hpf followed by PCR amplification (Wnt16 F: FAM –GCCTGGTTATGGCATTTCAA, Wnt16 R: AAAACAAAACGTAAATGTGAGACA) and fragment length analysis (ABI 3500) (Fig. S9B), as described previously (Carrington et al., 2015). After selecting the most efficient gRNA, (>90% of injected embryos subjected to fragment analysis showed indel mutations in Wnt16 (e.g. Fig. S9B) n=45), injections were carried out in eggs from incrosses of Tg(Sox10:GAL4-VP16);Tg(UAS:Kaede) or Tg(Col2a1aBAC:mcherry) followed by Kaede photoconversion and imaging. Amira (version 6.3) was used to 3D render Tg(Col2a1aBAC:mcherry) jaw cartilage.

### Mouth movements

Zebrafish anaesthetised with MS222 were mounted laterally on coverslips in 1% agarose. Forceps were used to remove agarose from around the head and Danieau solution was repeatedly flushed over the agarose-free cavity around the head until normal jaw movements resumed. The number of jaw movements in 1 minute were recorded, using a stereo microscope.

### In situ hybridisation

*In situ hybridisation* was performed as described (Thisse and Thisse, 2008), using a *lef1 probe* on 3dpf larvae. *Lef1* plasmid in a pBS-SK vector with Ampicillin resistance was delinearised using EcoRI and transcribed using T7. Samples were stored in 70% glycerol and wholemount larvae imaged using a stereo microscope or jaws were dissected and imaged using a compound microscope.

### Statistics

Statistics were performed using SPSS software. Student t-test and Mann-Whitney U test were used for comparisons between parametric and non-parametric data, respectively. One-way ANOVA and Kruskal–Wallis tests were used to make multi-comparisons between parametric and non-parametric data, respectively. The test for each experiment is reported in the figure legend.

## Data Availability

Raw data will be made available with a DOI through the data.bris.ac.uk server upon manuscript acceptance.

## Acknowledgements

We would like to thank members of the Wolfson Bioimaging facility and Dominic Alibhai for Multiphoton microscopy support, Robert Knight for plasmids, Karen Roddy for statistics advice and members of the Hammond lab for discussions during manuscript preparation.

